# Molecular epidemiology of mumps viruses detected in the Netherlands, 2017-2019

**DOI:** 10.1101/2020.04.30.070243

**Authors:** Rogier Bodewes, Linda Reijnen, Jeroen Kerkhof, Jeroen Cremer, Dennis Schmitz, Rob van Binnendijk, Irene K. Veldhuijzen

## Abstract

Mumps cases continue to occur, also in countries with a relatively high vaccination rate. The last major outbreaks of mumps in the Netherlands were from 2009-2012 and thereafter, only small clusters and single cases were reported. Molecular epidemiology can provide insights in the circulation of mumps viruses. The aims of the present study were to analyze the molecular epidemiology of mumps viruses in the Netherlands in 2017-2019 and to elucidate whether complete genome sequencing adds to the molecular resolution of mumps viruses when compared to sequencing of the mumps SH gene and non-coding regions (SH+NCRs). To this end, Sanger sequence data from the SH+NCRs were analyzed from 82 mumps genotype G viruses. In addition, the complete genomes were obtained from 10 mumps virus isolates using next-generation sequencing. Analysis of SH+NCRs of mumps viruses revealed the presence of two major lineages in the Netherlands, which was confirmed by analysis of complete genomes. Comparison of molecular resolution obtained with SH+NCRs and complete genomes clearly indicated that additional molecular resolution can be obtained by analyzing complete genomes. In conclusion, analysis of SH + NCRs sequence data from recent mumps genotype G viruses indicate that mumps viruses continue to circulate in the Netherlands and surrounding countries. However, to understand exact transmission trees and to compare mumps viruses on a large geographic scale, analysis of complete genomes is a very useful approach.

## Introduction

Mumps viruses are single stranded negative-sense RNA viruses belonging to the genus *Orthorubulavirus* of the family of *Paramyxoviridae*. Infection of humans with mumps virus results in acute illness which is usually characterized by a temporary unilateral or bilateral parotitis. Occasionally, mumps virus infections can result in serious complications [1, 2], but infections with mumps virus also often do not result in recognized clinical signs [1, 3].

In 1987, vaccination against mumps virus was implemented in the National Immunization Program in the Netherlands. Although this resulted in a rapid decline of mumps cases, in recent years multiple outbreaks occurred mainly among vaccinated young adults [4, 5]. From 2013 until 2020, multiple small local outbreaks and individual mumps cases were reported in the Netherlands.

Molecular epidemiology can provide insights in the circulation of mumps viruses. Sequencing of the SH gene and adjacent non-coding regions provides information about the genotype and some information of the circulating strains [6, 7]. Increasing the molecular resolution by obtaining additional sequence data of mumps viruses has proven useful in defining different transmission chains and local clusters [6, 8, 9]. In this context, sequencing of the hemagglutinin-neuraminidase protein gene (HN) and fusion protein gene (F) provides similar sequence resolution compared to sequencing of the sum of three the non-coding regions (NCRs), present between the nucleocapsid protein and phosphoprotein (N-P), between phosphoprotein and matrix protein (P-M) and between matrix protein gene and F gene (M-F). While both contributes to the molecular resolution, sequencing of the complete genome might provide the more complete resolution that is necessary to compare mumps viruses on a larger geographic scale and identify exact transmission chains [10-14].

The aims of the present study were to analyze the molecular epidemiology of mumps viruses in the Netherlands in 2017-2019 and to elucidate whether complete genome sequencing adds to the molecular resolution of mumps viruses when compared to direct sequencing of the mumps SH gene and NCRs. To this end, we investigated all typable specimens generated from the isolated viruses of molecular confirmed mumps cases in the Netherlands from 2017 to 2019.

## Materials and methods

### Sample collection and Sanger sequencing

Clinical samples were obtained from mumps cases in the Netherlands that were notified under the Public Health Act in the Netherlands. Samples were send to the National Institute for Public Health and the Environment (RIVM) for molecular surveillance. Sequencing of the SH region and the non-coding regions (NCRs) between the N-P, P-M and M-F genes was performed directly on clinical materials as described previously [6, 7].

### Virus isolation

In total 73 clinical materials (oral fluid, throat swabs and urine) were used for virus isolation. Confluent Vero cells (ECACC 84113001) in 24 wells plates (Greiner bio-one) were inoculated with clinical material (5μl oral fluid, 10µl urine or 50μl throat swab medium) and cells were incubated for one hour at 37°C/5% CO_2_. Subsequently, 1 ml of Dulbecco’s Modified Eagle Medium (DMEM, Gibco) was added supplemented with 100U/ml Pen-Strep (Lonza), 2 mM L-Glutamine (Lonza), 12.5 μg/ml Amphotericin B (Biowest) and 24 units/ml Nystatin suspension (Sigma) and plates were incubated at 37°C/5% CO_2._ Wells were checked daily for the presence of cytopathic effect (CPE). If no CPE was present after 5 days, cells were trypsinized (0.25% Trypsin-EDTA(1X) Gibco) and subsequently about 1/3 of trypsinized cells were transferred to another well and fresh medium was added. Again, cells were checked daily for the presence of CPE. If no CPE was present for 12 days after the start of the first incubation, culture was considered negative. If CPE was detected, medium was harvested and subsequently passaged one or two times over confluent Vero cells to produce large virus stocks (T25 cm^2^ flask or T75 cm^2^ flask, Corning).

### Next-generation sequencing

Mumps virus isolates were processed for full genome sequencing using a viral metagenomics approach. To this end, 200µl of each cell culture supernatant was pretreated and processed for next-generation sequencing using the Illumina NextSeq 550 platform essentially as described elsewhere [15](Benschop et al, manuscript in prep). Bioinformatic analysis of obtained reads was performed using the Jovian pipeline (Schmitz *et al*, manuscript in prep).

### Genetic and phylogenetic analysis

Obtained sequence data were aligned manually in MEGA7 and phylogenetic analysis was performed using the maximum-likelihood method with the Hasegawa-Kishino-Yano model and 500 bootstrap replicates [16]. Multiple closely related or mumps virus strains were added to the alignments for comparison.

## Results

### Mumps cases in 2017, 2018 and 2019 in the Netherlands

Between January 2017 and October 2019, 246 mumps cases were notified in the Netherlands based on clinical symptoms and laboratory confirmation, or an epidemiological link with a laboratory confirmed case. A mumps virus genotype could be determined for 123 cases (50%), of which 108 cases (88%) belonged to genotype G. The other 15 cases belonged to either genotype C (five cases), D (one case), H (seven cases), J (one case) or K (one case). From in total 82 mumps genotype G cases could NCRs sequence data be obtained in addition to the SH sequence used for genotyping (Genbank accession numbers MT238691-MT238955).

### Virus isolation

Inoculation of 9 throat swabs and one oral fluid specimen (14%) resulted in a mumps virus isolate as determined by the presence of cytopathic effect. Eight mumps virus isolates belonged to genotype G, while the other two mumps virus isolates belonged to genotype C or K (**Table 1**). Near complete genomes were obtained from all isolates, with genome sizes ranging from 15144 to 15375 nucleotides (Genbank accession numbers MT238681-MT238690).

**Table 1.** Overview of metadata of mumps virus isolates

| Isolate | Clinical specimen | Epidemiological link | Genotype | Mean read coverage |
| --- | --- | --- | --- | --- |
| MuVi/Amsterdam.NLD/29.17/1 | Throat swab | Southern Europe, travelling. | G | 3200* |
| MuVi/Beugen.NLD/16.18 | Throat swab | The Netherlands, unknown. | G | 4300 |
| MuVi/Eindhoven.NLD/42.18 | Oral fluid | The Netherlands, cluster among students at a university. | G | 550 |
| MuVi/Utrecht.NLD/8.19 | Throat swab | Southeast Asia, travelling. | K | 2000 |
| MuVi/Noord-Holland.NLD/9.19 | Throat swab | The Netherlands, unknown. | G | 2500 |
| MuVi/Deventer.NLD/12.19 | Throat swab | The Netherlands, cluster among students at a university. | G | 2300 |
| MuVi/Utrecht.NLD/15.19 | Throat swab | The Netherlands, contact with students at an university, possibly same cluster as MuVi/Deventer.NLD/12.19. | G | 480 |
| MuVi/Utrecht.NLD/21.19 | Throat swab | East-Asia, travelling. | C | 1400 |
| MuVi/Gelderland.NLD/26.19 | Throat swab | The Netherlands, work. | G | 3700 |
| MuVi/Utrecht.NLD/35.19 | Throat swab | The Netherlands, unknown. | G | 2600 |

\*if the sequence consisted of two or more overlapping contigs by de-novo assembly, the mean coverage of the contig with the lowest coverage is indicated.

### Comparison of sequencing data of isolates with original materials

Comparison of nucleotide sequence data of the SH gene and adjacent NCRs and N-P, P-M and M-F NCRs obtained from mumps viruses detected in clinical materials by Sanger sequencing with the consensus sequence data from isolates obtained by NGS revealed in total three differences. At position 194 of the SH gene of MuVi/Deventer.NLD/12.19 a C was present, while a T was detected at that position in the original material of MuVs/Deventer.NLD/12.19. Sanger sequencing confirmed the presence of a T at this position in the original clinical sample. Additional Sanger sequencing revealed the nucleotide shift from T to C was already present after the first passage.

At position 442 of the N-P NCR, a C was detected in the sequence data from MuVi/Amsterdam.NLD/29.17/1, while an A was detected in the sequence data obtained from the original material (MuVs/Amsterdam.NLD/29.17/1). Since this nucleotide position was in the phosphoprotein, this nucleotide change resulted also in an amino acid change (lysine to glutamine). The presence of the A in the original material was confirmed by Sanger sequencing. Additional Sanger sequencing analyses of the various passages indicated that the C was already detected at this position as a minor variant after the first passage, while two peaks of equal size were detected after the second passage. In the P-M NCR of MuVs/Utrecht.NLD/35.19, two peaks of similar size (A/G) were detected in the Sanger sequence data (both forward and reverse sequence), while the consensus sequence of MuVi/Utrecht.NLD/35.19 was a G at that position. This nucleotide position was also in the phosphoprotein, but this nucleotide difference did not result in an amino acid change. Additional analysis of the first passage and second passage of this isolate by Sanger sequencing indicated that in all these passages both nucleotide variants were present with two peaks of equal size. No other nucleotide differences were detected between the sequence data obtained from the original materials and the isolates.

### Phylogenetic analysis of concatenated SH+NCRs sequences

Phylogenetic analysis of concatenated SH+NCRs mumps genotype G virus sequences (Genbank accession numbers indicated that two main lineages of genotype G virus strains were present in the Netherlands in 2017-2019 with respectively 50 and 17 mumps viruses (**Figure 1**), which differed by in total 8 nucleotides in the SH and NCRs. Viruses of the first lineage were only detected in the second half of 2018 and 2019, while viruses from the second lineage were detected in 2017, 2018 and 2019. Another lineage of mumps genotype G viruses consisted of 6 cases but within this group more sequence variation was present (lineage 3). Within these three lineages, multiple sequence variants were detected in the Netherlands, of which a number belonged to known epidemiological clusters. In addition, various other virus variants were detected in the Netherlands in 2017, 2018 and 2019, but these variants were only detected in a limited amount of cases. Eight out of 15 of these viruses were collected from persons that were infected abroad. The presence of two main lineages was confirmed by analysis of (near) complete genomes of mumps viruses detected in these cases (**Figure 2**).

**Figure 1.**
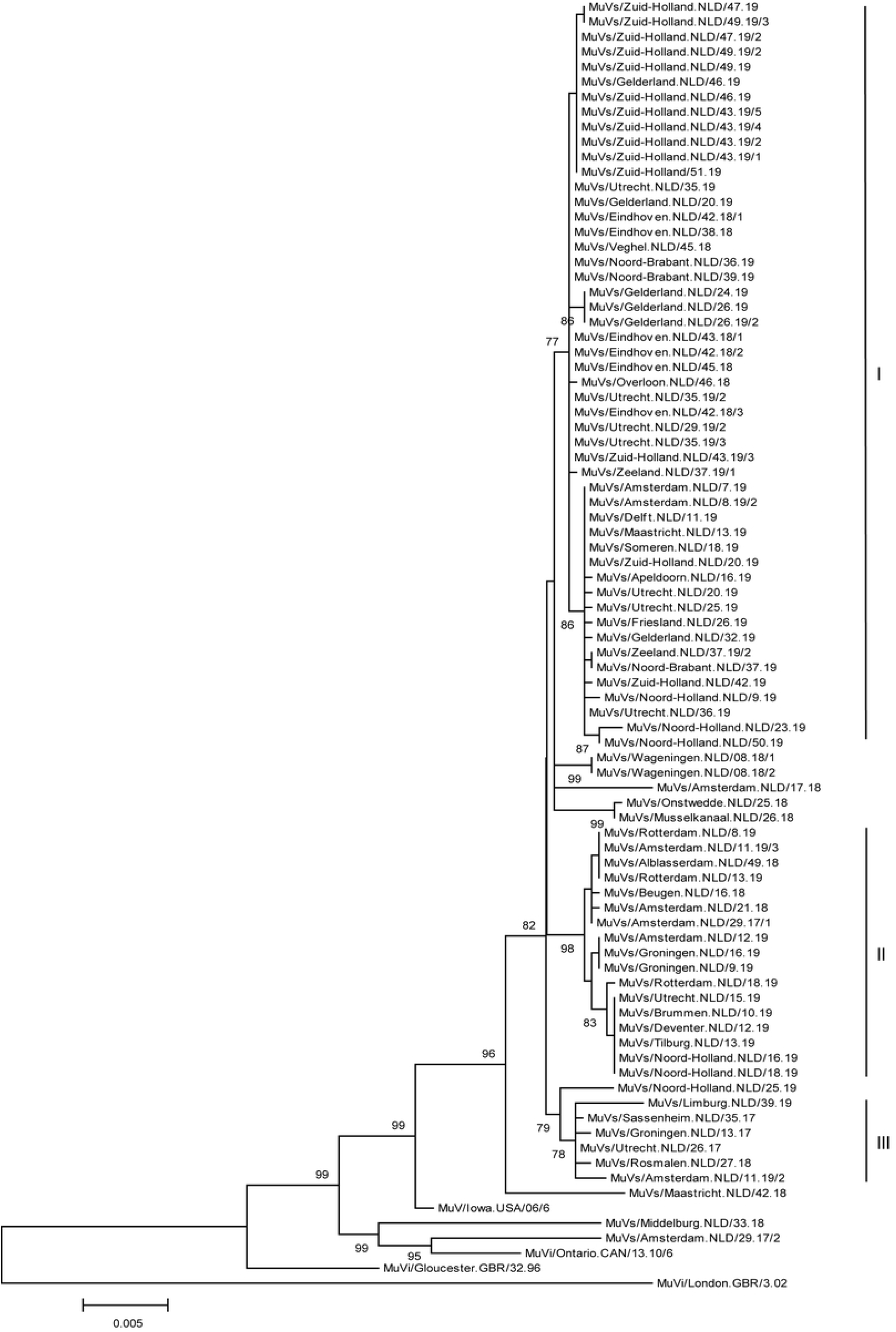
Phylogenetic analysis of concatenated sequence data of SH and NCRs of mumps viruses detected in the Netherlands from 2017-2019. Only bootstrap values >70 are indicated. The main lineages of mumps viruses based on phylogenetic analysis are indicated with I, II and III.

**Figure 2.**
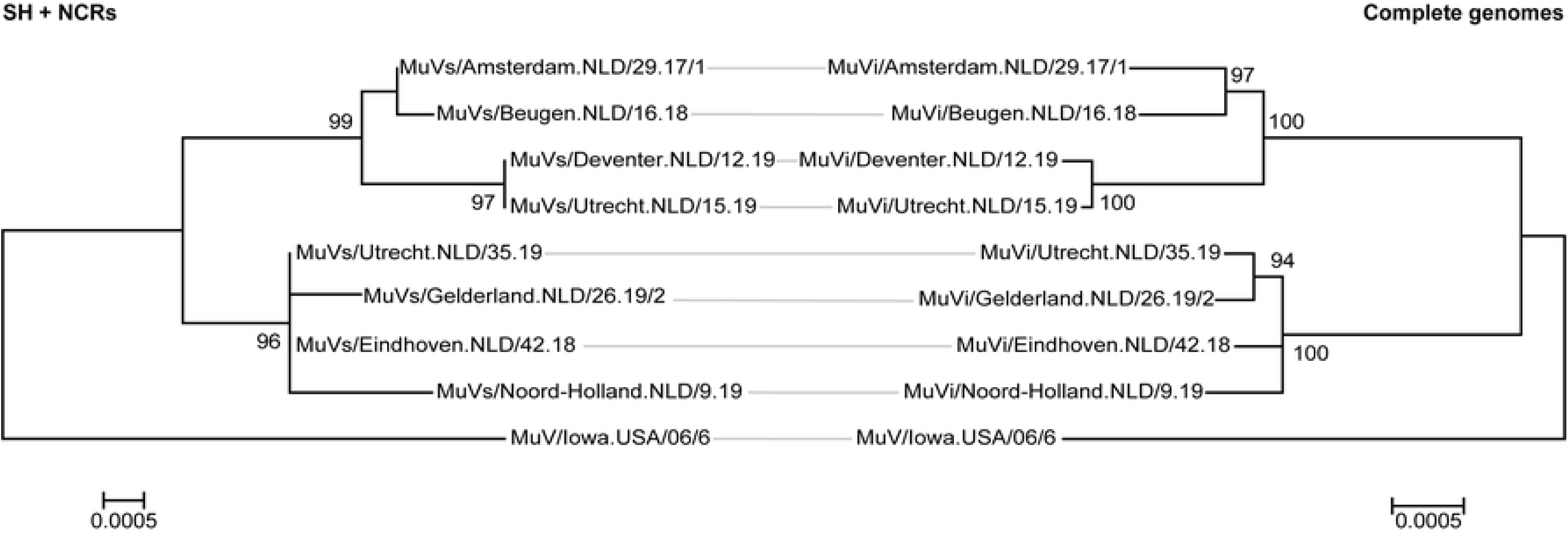
Comparison of molecular resolution obtained by analysis of SH and NCRs and complete genomes. Phylogenetic analysis was performed on complete genomes of isolates of mumps genotype G viruses and results were compared with SH+NCRs data obtained from the same set of mumps viruses detected in clinical materials. Please note that one nucleotide difference was detected in mumps virus isolates MuVi/Deventer.NLD/12.19, MuVi/Amsterdam.NLD/29.17/1 and MuVi/Utrecht.NLD/35.19 in the SH and NCRs compared to the sequences from mumps viruses detected in the clinical materials. Only bootstrap values >70 are indicated.

### Comparison of molecular resolution of SH+NCRs and complete genomes

Comparison of the molecular resolution provided by concatenated SH+NCRs sequences and (near) complete genomes revealed that there was an increase in molecular resolution if (near) complete genomes were analyzed.. For example, one nucleotide difference was present between the SH+NCRs sequences of MuVi/Amsterdam.NLD/29.17/1 and MuVi/Beugen.NLD/16.18, while 16 nucleotide differences were present between the isolated complete genomes of these two viruses. Furthermore, MuVs/Deventer.NLD/12.19 and MuVs/Utrecht.NLD/15.19 were identical for SH+NCRs sequences, but four nucleotide differences were present between the complete genomes of these isolated viruses. Also SH+NCRs sequences of MuVs/Utrecht.NLD/35.19 and MuVs/Eindhoven.NLD/42.18/1 were identical, while 11 nucleotide differences were detected by analysis of complete genomes. Although similar branching was observed in phylogenetic trees prepared by analysis of concatenated SH+NCRs and complete genomes, some differences were detected between the positions of MuVs/Gelderland.NLD/26.19/2 and MuVs/Noord-Holland.NLD/9.19 in the SH+NCR tree and the complete genome tree (**Figure 2**).

### Phylogenetic analysis of near complete mumps viruses

Pairwise identity analysis on the nucleotide level of mumps genotype G viruses of the complete genomes of the MuVi/Gelderland.NLD/26.19/2, MuVi/Utrecht.NLD/35.19, MuVi/Eindhoven.NLD/42.18 and MuVi/Noord-Holland.NLD/9.19 indicated that these viruses were most closely related to MuVs/Michigan.USA/4.16 (99.91%). MuVi/Amsterdam.NLD/29.17/1, MuVi/Beugen.NLD/16.18, MuVi/Deventer.NLD/12.19 and MuVi/Utrecht.NLD.15.19 were most closely related to MuVs/Montana.USA/11.16 and MuVs/Illinois.USA/26.15/2 (99.63%). In addition, genotype K virus MuVi/Utrecht.NLD/8.19 was most closely related to MuVs/Massachusetts.USA/24.17/5[K] (99.37%) and genotype C virus MuVi/Utrecht.NLD/21.19 was most closely related to MuVi/Kushinagar.IND/36/13[C] (99.01%) (**Figure 3**).

**Figure 3.**
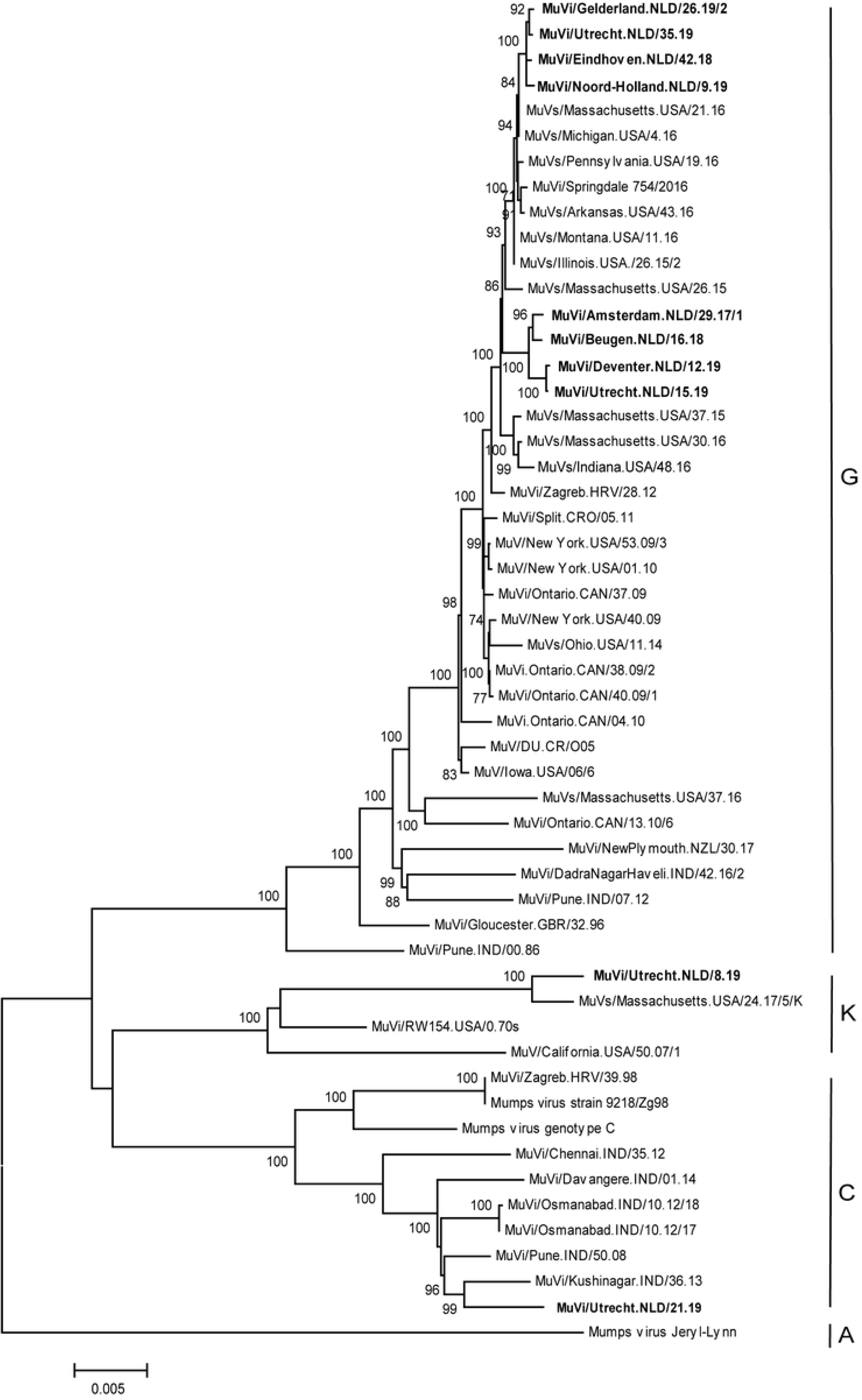
Phylogenetic analysis of complete mumps virus genomes. Phylogenetic analysis was performed on complete genomes of mumps virus isolated detected in the Netherlands and various closely related and representative mumps viruses with the HKY model and 100 bootstrap replicates. Only bootstrap values >70 are indicated. Mumps viruses detected in the Netherlands are indicated in bold. Genotypes to which these viruses belong are indicated.

## Discussion

Mumps cases continue to occur, also in countries with a relatively high vaccination rate. While in various countries recently large outbreaks were reported among young adults, the last major outbreaks of mumps in the Netherlands were from 2009-2012. Thereafter, only small clusters and single cases were reported [17].

Use of molecular epidemiology could contribute to the understanding of mumps virus circulation, also in countries or regions without major outbreaks. Therefore, the molecular epidemiology of mumps virus was studied using SH+NCRs mumps virus sequences obtained from mumps cases in the Netherlands in 2017-2019. Analysis of these regions with the relatively highest variation of mumps viruses revealed the presence of two major lineages in the Netherlands. The presence of these two lineages was confirmed by analysis of near complete genomes of a selection of these mumps viruses.

The presence of two lineages with multiple closely related mumps viruses in recent years, suggests that there was endemic transmission of mumps in the Netherlands and surrounding countries. This is supported by the fact that complete genomes of both lineages are distinct from mumps viruses detected recently in mainly the United States of America and Canada [10, 18]. Of interest, viruses of the first lineage were not detected before the second part of 2018, suggesting that this lineage emerged recently in the Netherlands.

In addition to viruses from these two lineages, multiple viruses were detected that had several nucleotide differences compared to viruses from these two lineages. Most likely, these viruses are currently not circulating in the Netherlands and surrounding countries, but were collected from solitary import cases or small clusters after initial import. This is supported in most cases by epidemiological data. Furthermore, mumps viruses were detected with other mumps genotypes than genotype G. Since these mumps viruses were detected only in a relatively small proportion, these viruses were most likely not endemic in the Netherlands in recent years.

Of interest, comparison of NCR+SH sequence data from original materials with that of isolates indicated that passaging of mumps viruses over cells resulted in three nucleotide changes in total. Since the sequence of 2270 nucleotides of in 10 total viruses were compared, it was calculated that 0.013% of all nucleotide positions changed due to passaging (∼2 nucleotide changes for a full mumps virus genome). Although analysis of these partial genomes indicate that passaging of mumps viruses results in nucleotide changes, this indicates that there is no major concern to use isolates for molecular surveillance studies in general. However, sequence data from isolates should be interpreted carefully if used to understand exact transmission trees. To study transmission trees, preferably original materials or eventually viruses that are passaged only a single time over cells are used.

Comparison of molecular resolution obtained with SH+NCRs and complete genomes clearly indicated that additional molecular resolution can be obtained by analyzing complete genomes. Main mumps clusters in recent years in the Netherlands could be identified by analysis of SH+NCRs data. However, NCR+SH sequence data cannot be used to study exact transmission trees, which is possible for mumps virus if complete genomes are analyzed [10, 18].

Most mumps viruses from which a complete genome was obtained in this study were collected from mumps cases that occurred several months after each other, and therefore it cannot be concluded whether these viruses are part of the same transmission chain. However, mumps viruses MuVi/Noord-Holland.NLD/9.19, MuVi/Deventer.NLD/12.19 and MuVi/Utrecht.NLD.15.19 were collected within 6 weeks. While the sequence of MuVi/Noord-Holland.NLD/9.19 was relatively distinct from the other two viruses, between MuVi/Utrecht.NLD.15.19 and MuVi/Deventer.NLD/12.19 were only 4 nucleotide differences present. Since both mumps cases also lived in the same city, they were most likely part of the same mumps cluster, but there was no direct epidemiological link between these two cases.

In conclusion, analysis of SH + NCRs sequence data from recent mumps genotype G viruses indicate that mumps viruses continue to circulate in the Netherlands and surrounding countries. However, to understand exact transmission trees and to compare mumps viruses on a large geographic scale, analysis of complete genomes is a very useful approach.

## Acknowledgements

The authors wish to thank medical microbiological laboratories and municipal health services for collecting and submitting clinical specimens that are used in this study and Tim Severs and Harry Vennema for setting-up NGS protocols.

